## Supplementary material for "Conditional Success of Adaptive Therapy: The Role of Treatment-Holiday Thresholds and Non-Existence of Optimal Strategies Revealed by Mathematical Modelling and Optimal Control": Supplemetary text and figures

### A. Details of the dependence of TTP on various parameters in neutral competition condition

By varying the initial proportion of resistant cells and the initial total cell count under different treatment-holiday thresholds (Figure S2A1-A3), we observed that when the initial total cell count is small and the initial proportion of resistant cells is high, the TTP is short. In this case, although the tumor size is small, it contains a large proportion of resistant cells, making it difficult for sensitive cells to suppress resistant cells, and thus cancer becomes harder to treat. As the initial total cell count increases and the initial proportion of resistant cells decreases, the TTP gradually increases. At this stage, although the number of resistant cells is low, the tumor is larger, requiring multiple treatment cycles in adaptive therapy to suppress resistant cells. Overall, larger adaptive thresholds, higher initial total cell counts, and lower initial proportions of resistant cells result in better treatment outcomes.

By varying the treatment-holiday threshold for dose-skipping and the initial proportion of resistant cells under different initial total cell proportions (Figure S2B1-B3), we found that lower treatment-holiday thresholds and higher initial proportions of resistant cells result in shorter TTPs. This is because tumors containing a high proportion of resistant cells make it difficult for adaptive therapy to reduce the total cell count below the threshold. As the adaptive threshold increases and the initial proportion of resistant cells decreases, the TTP gradually increases. When the adaptive threshold is high and the initial proportion of resistant cells is low, the TTP is maximized. At this stage, the tumor contains a significant number of sensitive cells, and a high threshold adaptive therapy can increase the number of treatment cycles, fully utilizing sensitive cells to suppress resistant cell growth. Overall, higher initial total cell counts are associated with longer TTPs.

By varying the treatment-holiday threshold for dose modulation and the initial total cell count under different initial proportions of resistant cells (Figure S2C1-C3), we observed that lower treatment-holiday thresholds and lower initial total cell counts result in shorter TTPs. This occurs because, despite the smaller tumor size, the low adaptive threshold leads to prolonged treatment cycles, causing significant damage to sensitive cells and reducing their ability to suppress resistant cells, resulting in shorter TTPs. As the adaptive threshold increases and the initial total cell count rises, the TTP gradually increases. When the adaptive threshold and the initial total cell count are both high, the TTP is maximized. At this stage, although the tumor size is large, the shorter treatment cycles allow sensitive cells sufficient time to recover and suppress resistant cell growth, resulting in longer TTPs. Overall, lower initial proportions of resistant cells are associated with longer TTPs.

Under the same conditions, simulations that varied the growth rates of sensitive and resistant cells revealed that when the ratio of the growth rate of sensitive cells to resistant cells ($r_S/r_R$) is greater than 1, indicating that the growth rate of sensitive cells exceeds that of resistant cells, the sensitive cells exhibit stronger competitiveness, resulting in longer TTPs. Conversely, when the ratio is less than 1, indicating that the growth rate of resistant cells exceeds that of sensitive cells, the TTP is shorter, and the higher the growth rate of resistant cells, the shorter the TTP. This relationship is illustrated in Figures S3 through S7.

### B. Details of comparison of adaptive therapy with other strategies in the weak competition condition

To investigate the probability of selecting adaptive therapy under varying parameters, we randomly varied five key model parameters from Table 1, creating a cohort of 1200 virtual patients. Simulations across different initial tumor burdens revealed that as the initial cell count increases, the probability of selecting adaptive therapy also increases (Main Text Figure 5D1-D7). However, when the competition coefficients satisfy $\alpha > 1$ and $\beta > 1$, and the initial cell count approaches the carrying capacity, the probability of selecting adaptive therapy decreases (Main Text Figure 5D8). This is because, under high competition coefficients, tumor competition becomes intense, and both MTD and adaptive therapy can control tumor size indefinitely, leading to a lower probability of selecting adaptive therapy with adaptive therapy (Figure S11).

In the case of conditional-improve, the differences in treatment outcomes among the four optimal strategies are small (approximately one month; $C_{\mathrm{TH0}} = 0.98$, $(\delta_1, \delta_2, C_{\mathrm{TH2}} - 1, C_{\mathrm{TH1}} - 1) = (0.25, 0.25, 0.05, 0.07)$,

$(T_D, T) = (9, 10)$). Since intermittent therapy requires a treatment holiday, the calculation of its optimal cycle revealed a one-day treatment holiday, resulting in treatment outcomes similar to MTD (Figure S8A). Considering cumulative drug toxicity, AT-S demonstrates the best efficiency among the four strategies (Figure S8B).

Subsequently, we varied the initial tumor parameters to analyze the treatment outcomes of the four strategies. When the initial proportion of resistant cells remains constant, and the initial cell count is small, the differences in TTP extension among the strategies are minimal. As the initial cell count increases, AT-S outperforms the other strategies in extending TTP, while AT-M shows the best Relative efficiency (Figure S8C).

When the initial tumor burden remains constant, and the initial proportion of resistant cells is small, AT-S outperforms the other strategies. However, as the initial proportion of resistant cells increases, MTD, AT-S, and IT outperform AT-M, with AT-S and AT-M demonstrating the best Relative efficiency among the four strategies (Figure S8D).

For the uniform-improve scenario, the differences in treatment outcomes among the four optimal strategies are minimal (approximately one month; $C_{\mathrm{TH0}} = 0.98$, $(\delta_1, \delta_2, C_{\mathrm{TH2}}-1, C_{\mathrm{TH1}}-1) = (0.25, 0.25, 0.05, 0.09)$, $(T_D, T) = (20, 21)$). Similar to the conditional-improve scenario, the optimal intermittent therapy cycle includes a one-day treatment holiday, resulting in treatment outcomes similar to MTD (Figure S9A). Considering cumulative drug toxicity, AT-S shows the best efficiency among the four strategies (Figure S9B).

When varying the initial total cell count while keeping the initial proportion of resistant cells constant, the differences in TTP extension among the four strategies are minor. MTD and IT outperform AT-S and AT-M in terms of TTP, but AT-S and AT-M have a better relative efficiency than MTD and IT (Figure S9C). When varying the initial proportion of resistant cells while keeping the initial tumor burden constant, MTD and IT outperform AT-S and AT-M in terms of TTP. However, AT-S and AT-M exhibit a better Relative efficiency compared to MTD and IT (Figure S9D).

For the uniform-decline scenario, the differences in TTP among the four optimal strategies are larger (approximately two months; $C_{\mathrm{TH0}} = 0.98$, $(\delta_1, \delta_2, C_{\mathrm{TH2}} - 1, C_{\mathrm{TH1}} - 1) = (0.25, 0.25, 0.05, 0.05)$, $(T_D, T) = (15, 18)$). Among the four strategies, AT-S and AT-M demonstrate the best treatment outcomes (Figure S10A). Regarding cumulative drug toxicity, AT-S achieves the best relative efficiency (Figure S10B).

When varying the initial total cell count while keeping the initial proportion of resistant cells constant, the differences in treatment outcomes are minor for small tumors. For larger tumors, AT-S shows the best treatment outcomes, while AT-M and AT-S have a better Relative efficiency than the other two strategies (Figure S10C). When varying the initial proportion of resistant cells while keeping the initial tumor burden constant, AT-S demonstrates the best treatment outcomes regardless of the proportion of resistant cells. AT-M shows slightly worse treatment outcomes than the other three strategies but exhibits the best Relative efficiency (Figure S10D).

#### C. Supplementary figures

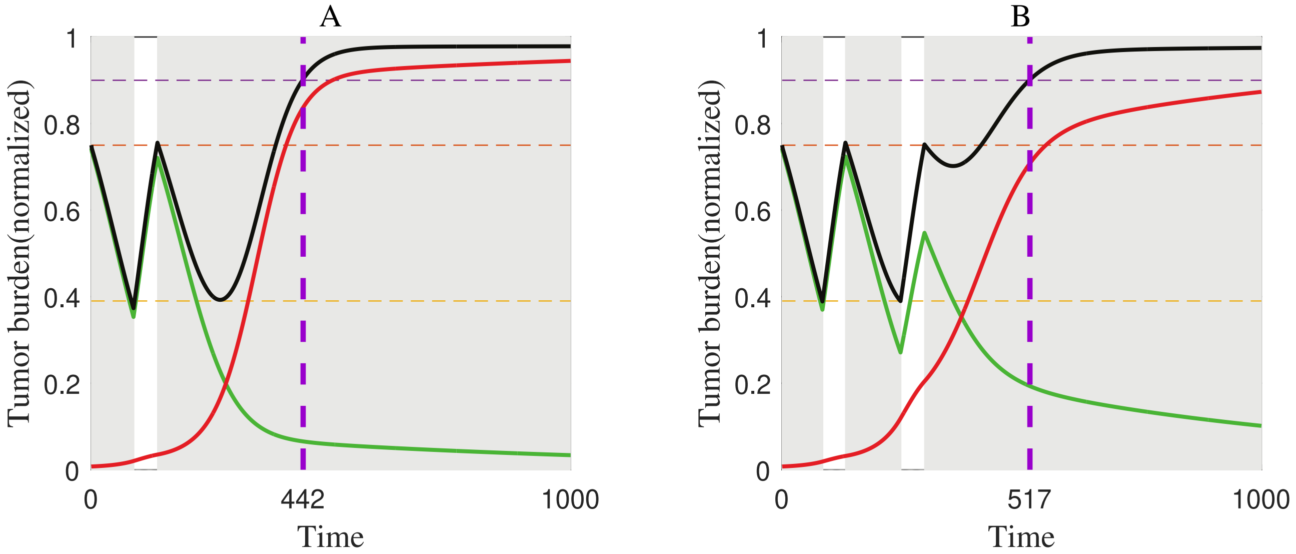


Figure S1. Tumor evolution dynamics in neutral competition with different treatment-holiday thresholds, showing the reason of discontinuity in the dependence of TTP on the treatment-holiday thresholds (Figure 2). Purple dashed line: TTP; Red: Resistant cells; Green: Sensitive cells; Black: Total cell count; Shaded area: Treatment implementation. **(A)** $n_0 = 0.75$, $f_R = 0.01$, $C_{\mathrm{TH0}} = 0.51$. **(B)** $n_0 = 0.75$, $f_R = 0.01$, $C_{\mathrm{TH0}} = 0.52$. Other parameters are taken as those in Table 1.

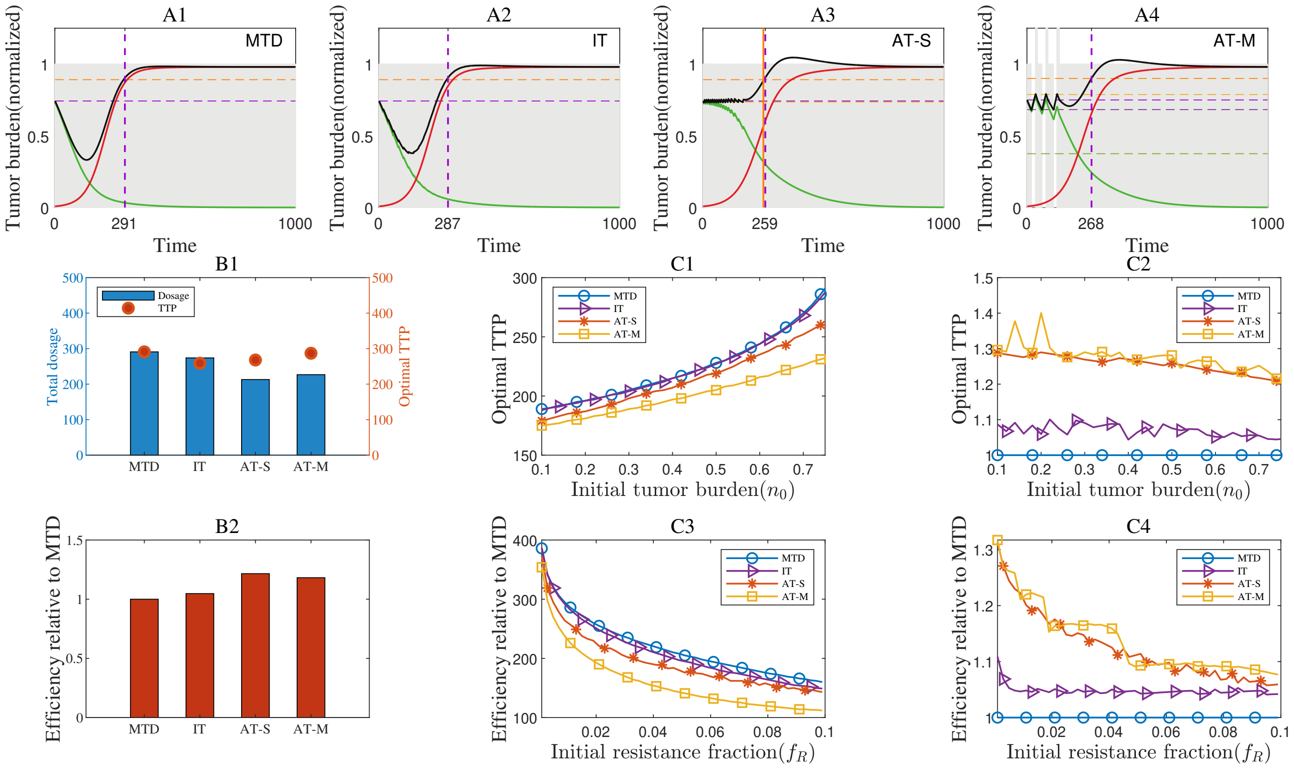


FIGURE S8. Comparison of four treatment strategies on TTP in the scenario of **conditional-improve** where the parameters are taken as $\alpha = 0.6$,$\beta = 0.5$, $f_R = 0.01$ and $n_0 = 0.75$. **(A1)-(A4)** Tumor dynamics under four treatment strategies with optimal treatment parameters. Adaptive therapy outperforms MTD and IT. Optimal treatment parameters: $C_{\text{TH0}} = 0.98$, $\delta_1 = 0.25$, $\delta_2 = 0.25$, $C_{\text{TH2}} = 1.05$ and $C_{\text{TH1}} = 1.07$, $T_D = 9$, $T = 10$. **(B1)** Cumulative drug toxicity (total dosage) and corresponding TTP for the four strategies. **(B2)** Relative efficiency for the four strategies. **(C1)-(C2)** Effects of varying the initial tumor cell proportion on TTP and Relative efficiency of the four strategies. **(C3)-(C4)** Effects of varying the initial resistant cell proportion on TTP and Relative efficiency of the four strategies.

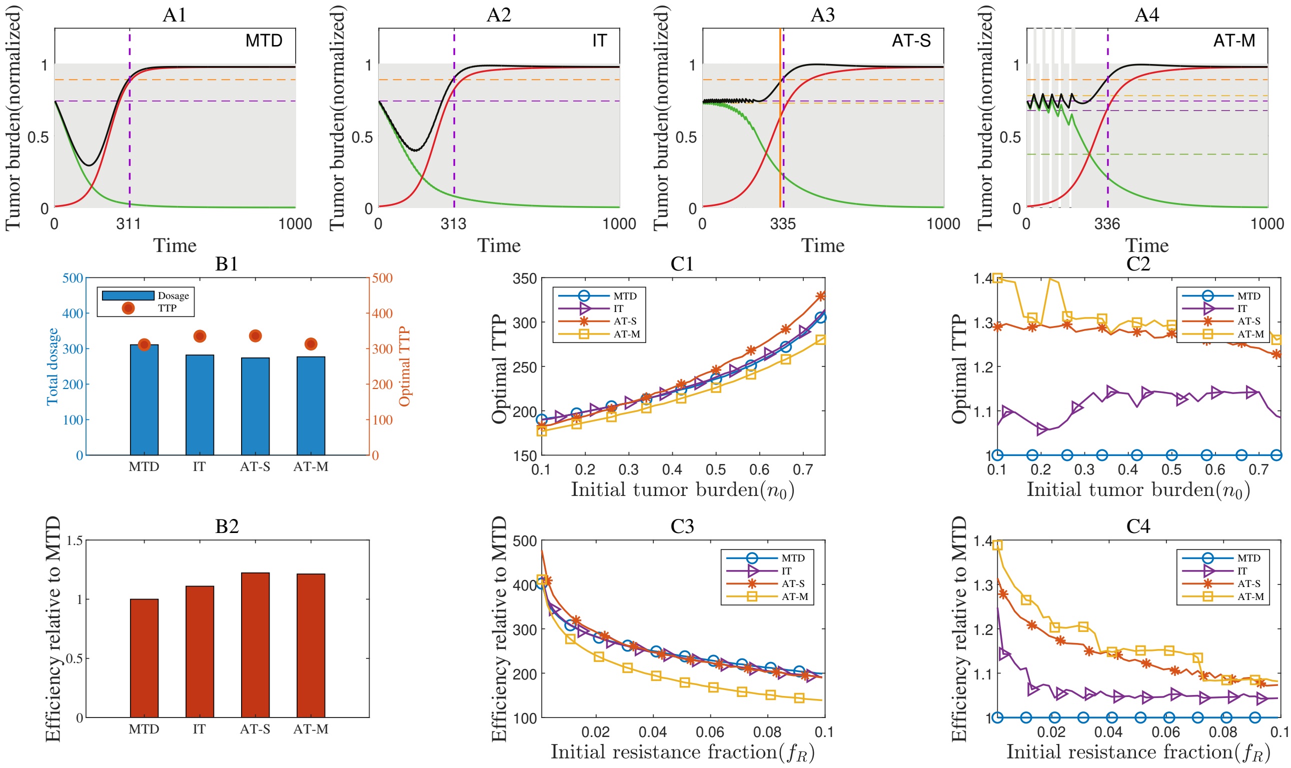


FIGURE S9. Comparison of four treatment strategies on TTP in the scenario of **uniform-improve** scenario where the parameters are taken as $\alpha = 0.6$, $\beta = 0.3$, $f_R = 0.01$ and $n_0 = 0.75$. **(A1)-(A4)** Tumor dynamics of tumor evolution with four treatment strategies with optimal treatment parameters. Optimal parameters are taken as follows. $C_{\text{TH0}} = 0.98$, $\delta_1 = 0.25$, $\delta_2 = 0.25$, $C_{\text{TH2}} = 1.05$ and $C_{\text{TH1}} = 1.09$, $T_D = 20$, $T = 21$. **(B1)** Cumulative drug toxicity (total dosage) and corresponding TTP for the four strategies. **(B2)** Relative efficiency for the four strategies. **(C1)-(C2)** Effects of varying the initial tumor cell proportion on TTP and Relative efficiency of the four strategies. **(C3)-(C4)** Effects of varying the initial resistant cell proportion on TTP and Relative efficiency of the four strategies.

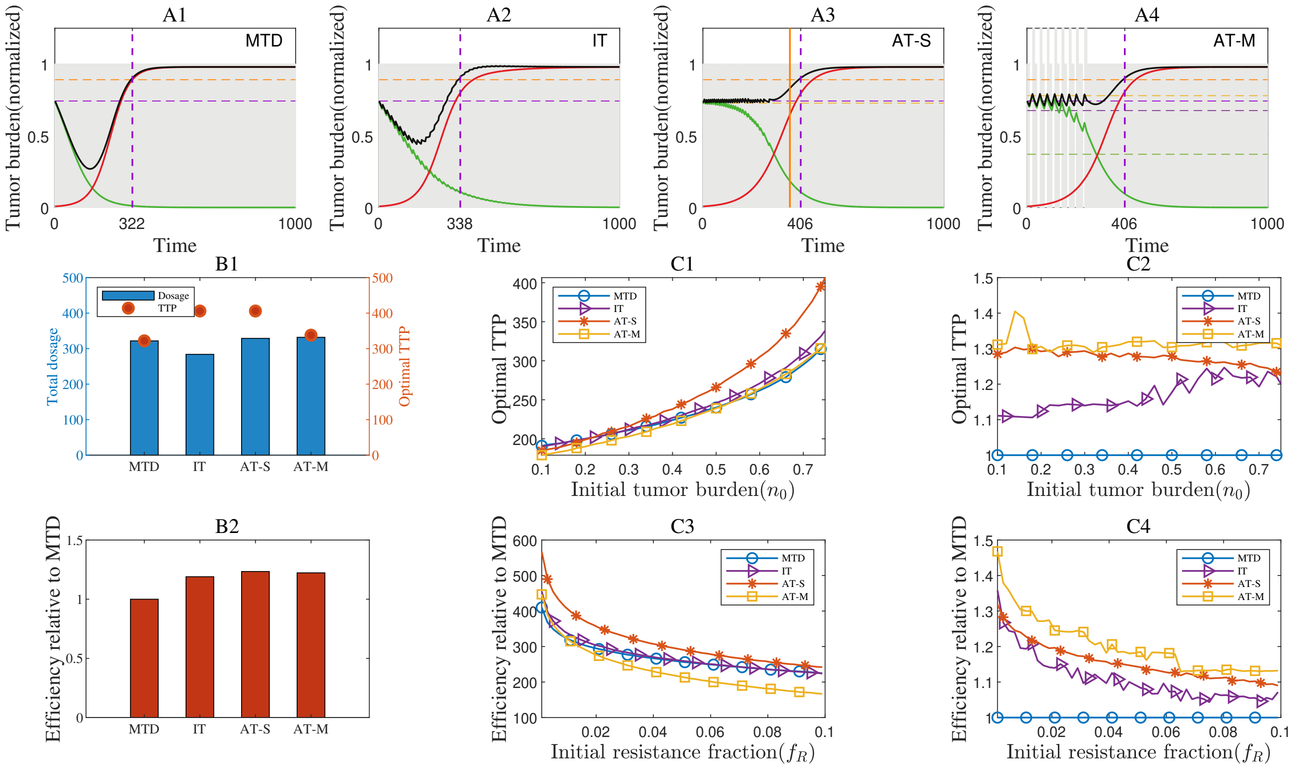


FIGURE S10. Comparison of four treatment strategies on TTP in the scenario of **uniform-decline** where the parameters are taken as $\alpha = 0.3$, $\beta = 0.6$, $f_R = 0.01$ and $n_0 = 0.75$) **(A)** Tumor dynamics under four treatment strategies with optimal treatment parameters. Optimal parameters are as follows. $C_{\text{TH0}} = 0.98$, $\delta_1 = 0.25$, $\delta_2 = 0.25$, $C_{\text{TH2}} = 1.05$ and $C_{\text{TH1}} = 1.05$, $T_D = 15$, $T = 18$. **(B1)** Cumulative drug toxicity (total dosage) and corresponding TTP for the four strategies. **(B2)** Relative efficiency for the four strategies. **(C1)-(C2)** Effects of varying the initial tumor cell proportion on TTP and Relative efficiency of the four strategies. **(C3)-(C4)** Effects of varying the initial resistant cell proportion on TTP and Relative efficiency of the four strategies.

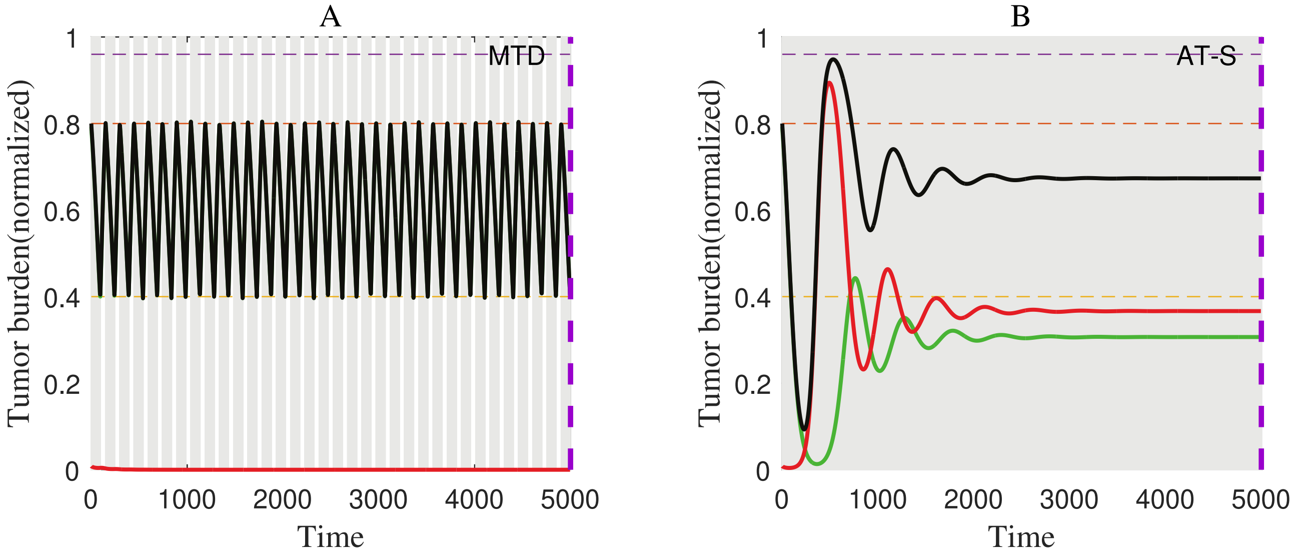


FIGURE S11. Tumor evolution dynamics with $\alpha = 2$, $\beta = 2$. Purple dashed line: TTP; Red: Resistant cells; Green: Sensitive cells; Black: Total cell count; Shaded area: Treatment implementation. **(A)** Adaptive therapy ($n_0 = 0.8$, $f_R = 0.01$, $C_{\text{TH0}} = 0.5$). **(B)** Maximum tolerated dose ($n_0 = 0.8$ , $f_R = 0.01$).
